# What shapes intraspecific genetic diversity in amphibians and reptiles? A comparative macrogenetic study across six biogeographic regions

**DOI:** 10.64898/2026.09.29.755451

**Authors:** Maria João Paúl, Matthew O. Moreira, André Vicente Liz, Sydney K. Decker, Craig Moritz, Bryan C. Carstens, Sílvia B. Carvalho

## Abstract

**Aim:** Given that much of Earth’s biodiversity is contained within species, inferring intraspecific genetic diversity and identifying its ecological correlates is essential for understanding evolutionary processes and informing conservation. Here, we investigate broad-scale variation in species-level mitochondrial genetic diversity and evaluate the relative importance of alternative ecological hypotheses proposed to explain it.

**Location:** Brazilian Atlantic Forest, Iberian Peninsula, northwestern Africa, Arabian Peninsula, Madagascar and Australian Monsoonal Tropics.

**Taxon:** Amphibians and reptiles

**Methods:** We assembled a comprehensive dataset by integrating and manually curating mitochondrial sequences, occurrence records, environmental data and species traits from multiple open-access sources. We used multiple mitochondrial genes to estimate mean nucleotide diversity for each study species and quantified regional variation in genetic diversity. We then used random forests models to evaluate the relative importance of selected variables representing hypotheses related to water-energy availability, topography, historical climatic stability, land-cover characteristics, species geographic characteristics, life-history traits, evolutionary history and conservation status.

**Results:** The final dataset comprised 247 amphibians and 295 reptiles, for which 43 and 52 explanatory variables were evaluated, respectively. Overall, the strongest support was found for hypotheses related to life-history traits, topographic heterogeneity and water-energy availability. In amphibians, higher genetic diversity was associated with smaller body size and greater environmental heterogeneity, particularly broader elevational and precipitation gradients. In reptiles, genetic diversity was primarily associated with life-history traits, with species of smaller body size and lower reproductive output exhibiting higher diversity. Long-term climatic stability showed comparatively little explanatory power for both taxa.

**Main conclusions:** Our comprehensive study demonstrates that integrating extensive manual data curation with comparative macrogenetic analyses provides a robust framework for identifying broad-scale ecological correlates of mitochondrial genetic diversity. These findings improve our understanding of the factors associated with contemporary patterns of genetic diversity and provide a foundation for future conservation assessments in data-limited systems.

## 1. Introduction

Global biodiversity is rapidly declining due to threats such as climate change and habitat loss (Convention on Biological Diversity, 2022; Peng et al., 2023). Maintaining genetic diversity within species is crucial to conservation since it is required for species to adapt in the face of rapid environmental change (Andrello et al., 2022; Sgrò et al., 2011). Intraspecific genetic diversity enhances community and ecosystem resilience and facilitates species survival and adaptation potential (Laikre et al., 2020). Understanding its spatio-temporal trends can also help to unveil the evolutionary and demographic processes that generate, maintain, and erode biodiversity (Leigh et al., 2021). As such, novel molecular techniques and computational tools have led to an exponential increase in the number of publicly available large-scale and high-quality genetic datasets (Blanchet et al., 2017). Macrogenetics has thus become a thriving field which aims at understanding broad-scale patterns of genetic diversity and their underlying drivers at large spatial, temporal and taxonomic scales (Blanchet et al., 2017; Leigh et al., 2021). These studies provide a remarkable opportunity to advance ecological and evolutionary theory and to improve long-term biodiversity conservation at multiple levels (Blanchet et al., 2017; Leigh et al., 2021; Schmidt et al., 2023).

A growing body of macrogenetic research proposed several environmental, geographic and biological hypotheses to explain variation in intraspecific genetic diversity (*e.g.,* Adams & Hadly, 2013; Amador et al., 2024; Doyle et al., 2015; Fonseca et al., 2023; Manel et al., 2020). These putative drivers primarily encompass hypotheses related to water and energy availability, historical climatic stability, topography, habitat configuration, species’ geographic range characteristics and biological attributes (Table 1). Although conceptually distinct and not mutually exclusive, these hypotheses predict that genetic diversity is shaped by a limited number of population genetic mechanisms. Climatic conditions determine energy and water availability, influencing mutation rates, demographic stability and effective population size, whereas topography and habitat configuration primarily regulate dispersal, connectivity and population subdivision. Likewise, species’ geographic characteristics, life-history traits, evolutionary history and conservation status are expected to be associated with demographic resilience, effective population size and the strength of genetic drift (De Kort et al., 2021; Lawrence et al., 2023). Collectively, these mechanisms influence the balance between mutation, genetic drift and gene flow, ultimately determining the amount and spatial configuration of genetic diversity maintained within species. Although many of these hypotheses derive from population genetic theory and studies of genome-wide nuclear diversity, they have been frequently explored using mitochondrial DNA (mtDNA), which remains the most widely available genetic marker for comparative macrogenetic studies.

**Table 1.**
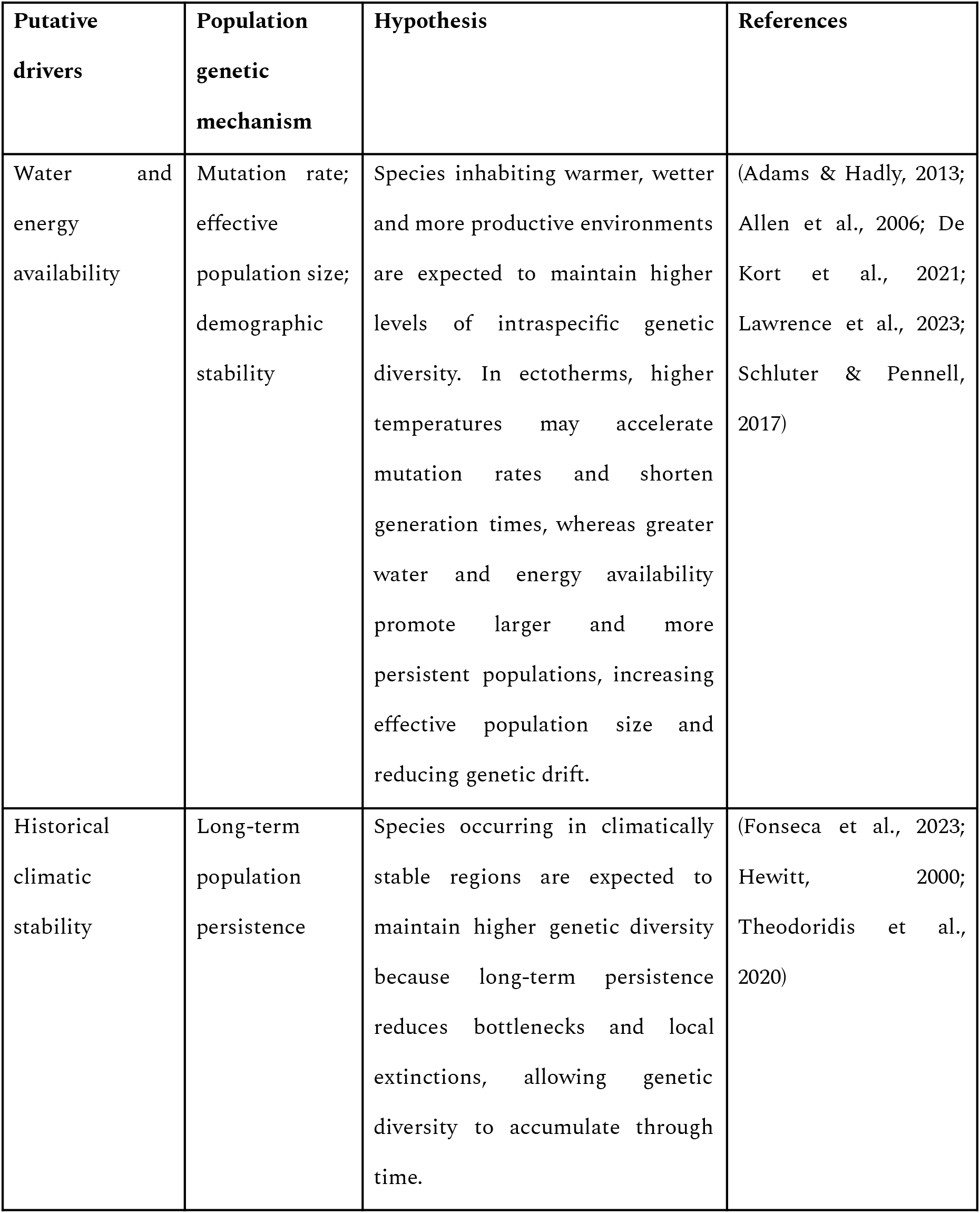
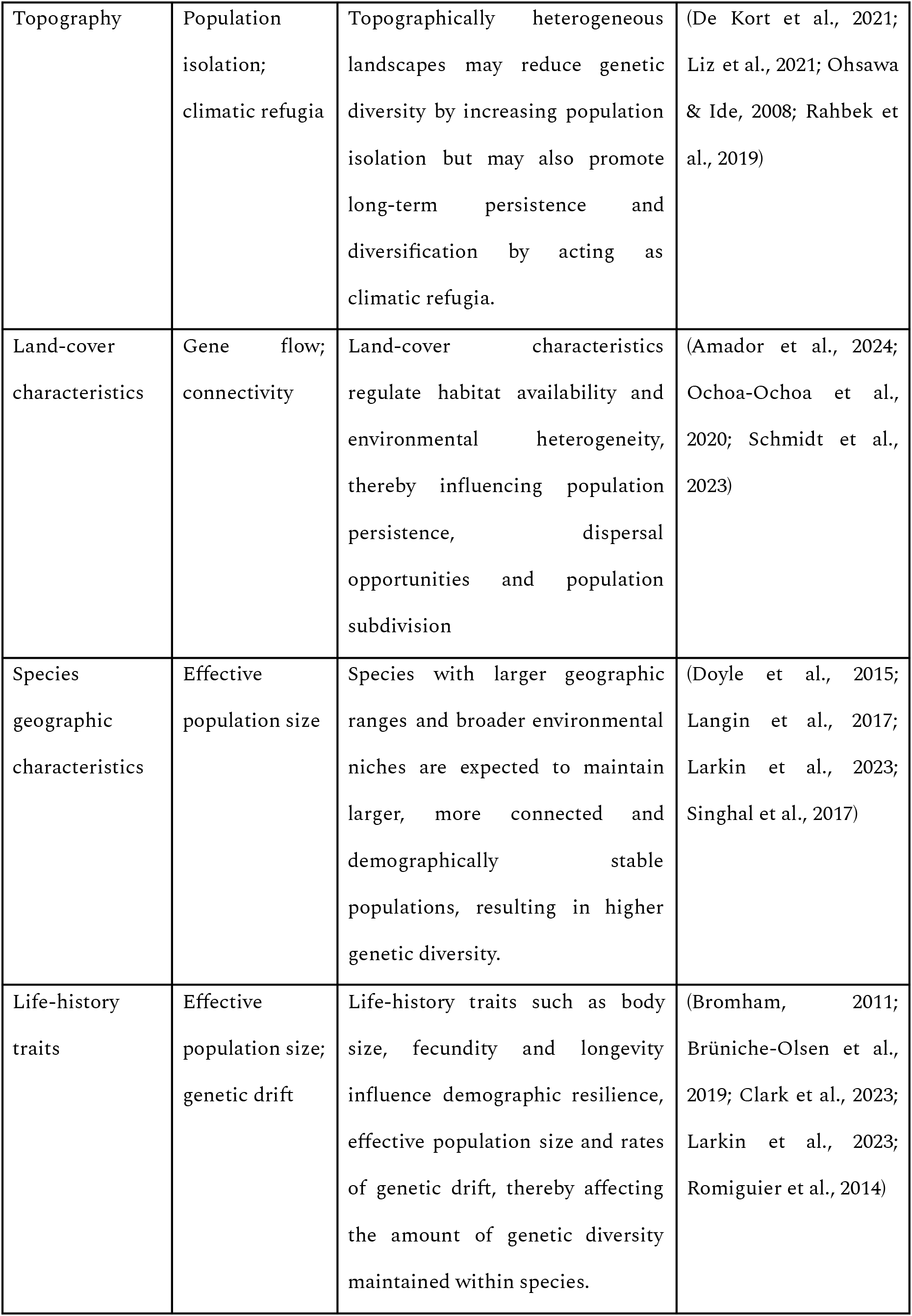
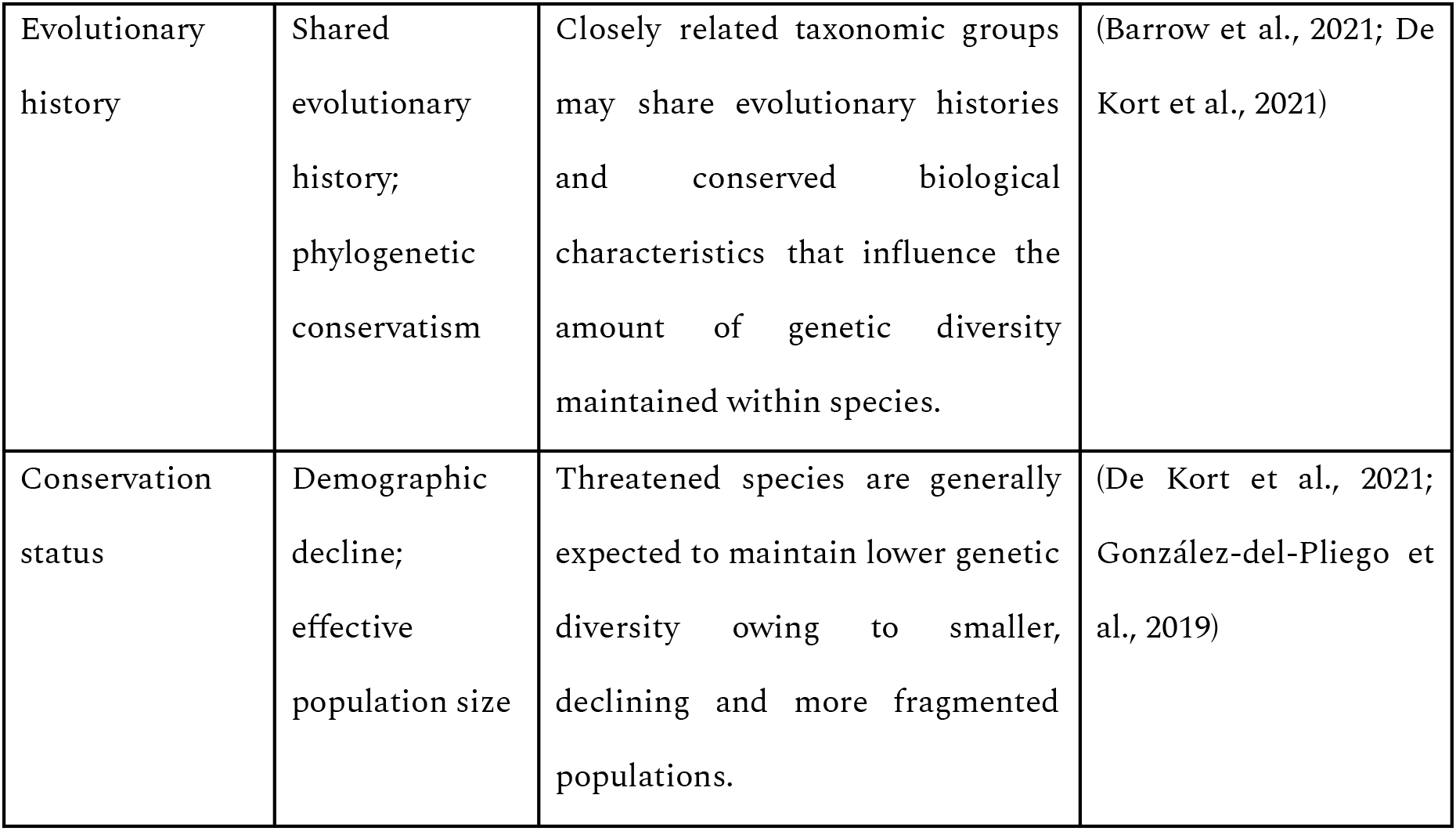
Summary of hypothesized predictors of genetic diversity in amphibians and reptiles evaluated in this study and their expected population genetic mechanism.

Despite the growing number of macrogenetic studies, it remains unclear which ecological hypotheses consistently explain variation in species-level genetic diversity because most studies evaluate only a subset of the potential environmental, geographic and biological mechanisms. Broad scale macrogenetic investigations often rely on machine learning algorithms to capitalise on large datasets and identify genetic diversity correlates (Amador et al., 2024; Barrow et al., 2021; Leigh et al., 2021) on automated computational approaches, which are prone to biases arising from incomplete geographic metadata, uneven spatial sampling and limited data curation (Leigh et al., 2021; Paz-Vinas et al., 2021). Indeed, most publicly available genetic data lack metadata describing sampling localities, limiting their reuse whenever spatial information is required (Pelletier et al., 2022). A complementary, yet underdeveloped, approach is to conduct macrogenetic studies on a representative regional scale, where it is possible to manually curate data to enhance dataset completeness through literature search, and to incorporate region-specific historical changes that might influence genetic diversity (*e.g.,* Amador et al., 2024).

Amphibians and reptiles represent an excellent model for macrogenetic research given their deep evolutionary histories, strong population structure and limited dispersal, as well as the abundance of publicly available genetic data generated by extensive phylogeographic research on these taxa. Furthermore, as ectotherms, their physiology, distribution and population dynamics are strongly influenced by climatic conditions making them exceptionally vulnerable to climate change (Foden et al., 2013). This can be especially concerning since past rapid climate change has been linked to declines in genetic diversity (Theodoridis et al., 2020), suggesting that historical fluctuations in climatic conditions have played a key role in shaping contemporary patterns of genetic variation.

In this study, we evaluate the relative importance of putative environmental, geographic and biological correlates on amphibian and reptile genetic diversity. The study spans six regions: Brazilian Atlantic Forest (hereafter Brazilian AF), Iberian Peninsula, northwestern Africa (hereafter NW Africa), Arabian Peninsula, Madagascar and Australian Monsoonal Tropics (hereafter AMT; Figure 1). These regions encompass a wide range of climatic conditions, environmental heterogeneity, geological histories and herpetofaunal communities. These regions also span tropical, Mediterranean, arid and monsoonal ecosystems, thereby maximizing variation in the ecological hypotheses evaluated in this study while allowing comparisons across contrasting evolutionary and environmental contexts. In addition, they have all been the focus of extensive phylogeographic research, providing unusually comprehensive mitochondrial datasets that permit rigorous manual curation and comparative analyses across hundreds of species.

**Figure 1.**
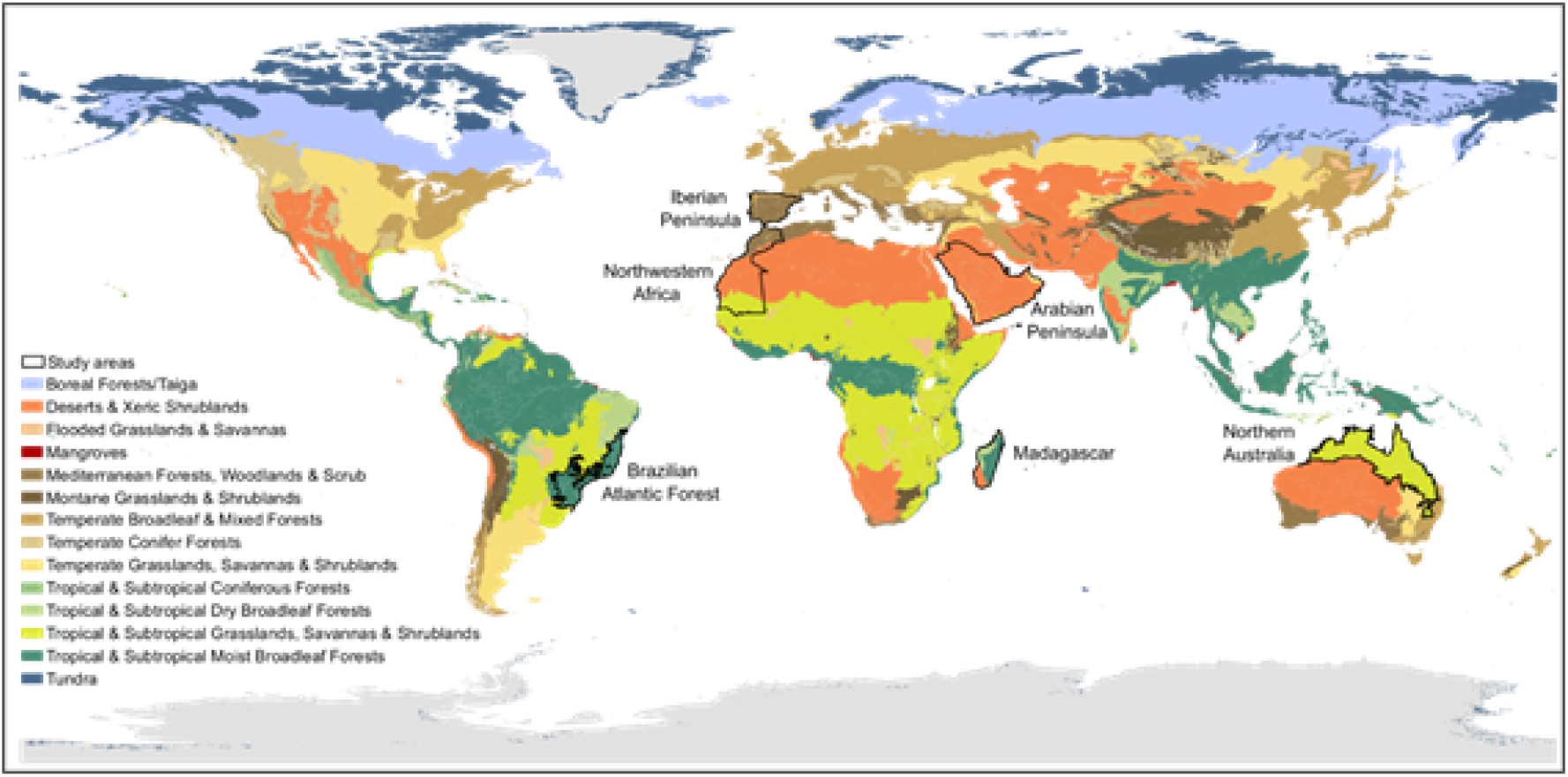
Six study regions and world biomes adapted from (Dinerstein et al., 2017).

This multi-regional framework provides an opportunity to investigate both large-scale geographic variation in genetic diversity and its potential correlates. Accordingly, we i) quantified variation in species-level intraspecific genetic diversity among study regions and ii) identified its main environmental, geographic and biological correlates using machine learning. Based on the hypotheses summarized in Table 1, we expect species-level genetic diversity to emerge from the combined influence of water-energy availability, topographic heterogeneity, historical climatic stability, habitat characteristics, species’ geographic attributes and life-history traits, rather than from any single population genetics mechanism. Because these mechanisms operate at different spatial and temporal scales, we further expect their relative importance to vary among biogeographic regions and between amphibians and reptiles.

In particular, we predict that species inhabiting regions characterized by higher precipitation, productivity, temperature, and climatic stability will maintain greater levels of genetic diversity, reflecting larger and more stable population sizes. Likewise, we expect species with broader niches, larger geographic ranges and life-history traits associated with larger effective population sizes to exhibit greater genetic diversity.

## 2. Material and methods

### 2.1. Compilation and curation of mitochondrial sequence data

We identified the species occurring in the six regions by intersecting all available amphibian and reptile range polygons from the IUCN Red List (IUCN, 2022) with the spatial boundaries of the six study regions (Figure 1). Then, for all herptile species occurring in the study regions, we obtained genetic sequences and associated metadata (including country, locality and coordinate information) from the NCBI repository (https://www.ncbi.nlm.nih.gov/genbank/) in a custom-built script using the rentrez R package (Winter, 2017). The full dataset targeted mtDNA sequence data, ranging from partial gene sequences to complete mitochondrial genomes. To reduce biases associated with small sample sizes, only species-marker combinations represented by at least ten sequences were retained for subsequent analyses (Barrow et al., 2021). We excluded marine and invasive species from our analysis because these are likely outliers in terms of the factors that influence genetic diversity.

To maximise spatial data compilation, we manually georeferenced sequences lacking NCBI spatial data by compiling coordinates and/or locality data from publications whenever such information was available. Then, for sequences that lacked coordinates but had locality data, we used the Google Maps API and the ggmap R package (Kahle & Wickham, 2013) to obtain approximate occurrences based on the locality descriptors filtered by country. We removed species without at least three georeferenced sequences for at least one mitochondrial marker.

For each marker, we performed a species-specific automated sequence alignment using SuperCRUNCH, a Python toolkit for assembling large phylogenetic datasets from NCBI sequences and/or unpublished data (Portik & Wiens, 2020). The overall workflow consisted of parsing the starting sequences, filtering and selecting sequences (we retained sequences with at least 250 bp), performing a multiple sequence alignment using the accurate adjust option in MAFFT, and conducting several post-alignment tasks such as relabelling sequence records for compatibility in downstream analyses and edge trimming to remove gap-rich regions from the alignments.

### 2.2. Estimation of species-level intraspecific genetic diversity

To estimate intraspecific genetic diversity for each species, we used a modified equation of nucleotide diversity (Nei & Li, 1979) as implemented in (Fonseca et al., 2023), which accounts for the different number of sequences present in our alignments. To maximise taxonomic representation while incorporating all available genetic information, we included every mitochondrial marker available for each species. Thus, for each mitochondrial marker, we used the nuc.div() function from the pegas R package (Paradis, 2010) to calculate nucleotide diversity between pairs of aligned sequences and then averaged it to obtain an estimation for each species and marker. Amphibian nucleotide diversity was log-transformed prior to statistical analyses to improve normality of the response variable and model residuals. Because mitochondrial markers differ in substitution rates and therefore in their expected levels of nucleotide diversity, we scaled herptile nucleotide diversity for each marker between zero and one. Then, for each species, we calculated mean nucleotide diversity of all available genes from the scaled values.

Differences in mean species-level genetic diversity among the six study regions were assessed using analysis of variance (ANOVA) implemented with the aov() function in the stats R package (R Core Team, 2024). Species occurring in multiple study regions were assigned to the region comprising the most of its range.

### 2.3. Compilation of environmental, geographic and biological predictors

To evaluate the hypotheses summarized in Table 1, we compiled a comprehensive set of environmental, geographic and biological variables representing each hypothesized driver of species-level intraspecific genetic diversity (Table 2). Specifically, we compiled variables describing climate, topography, land-cover characteristics, species’ geographic characteristics, ecological traits, taxonomy and conservation status. In addition, we quantified sampling effort and the spatial representativeness of genetic sampling, which were included as control variables to account for potential biases arising from uneven sequence availability and geographic sampling coverage (Table 2 and see Table S1).

**Table 2.** Environmental, geographic and biological variables compiled to represent the ecological hypotheses proposed in Table 1. Variables were obtained either directly from environmental layers, species range maps, taxonomic and trait databases, or derived from species distribution models (SDMs). Genetic sampling effort and spatial representativeness were included as control variables to account for potential sampling biases. Only variables selected through a recursive feature elimination analysis were included in the final models.

| Hypothesized driver | Compiled variables | Variable source | Description |
| --- | --- | --- | --- |
| Water and energy availability | Annual mean temperature (BIO1), isothermality (BIO3), maximum temperature of the warmest month (BIO5), minimum temperature of the coldest month (BIO6), annual precipitation (BIO12), precipitation of the wettest quarter (BIO16), precipitation of the driest quarter (BIO17) (range and sd); temperature niche breadth (SNBT), precipitation niche breadth (SNBP), climatic niche hypervolume, climatic niche marginality | Environmental layers; SDMs | Climatic variables representing energy and water availability, together with species' climatic niche characteristics, expected to influence mutation rates, demographic stability and effective population size. |
| Historical climatic stability | Climate Stability Index (CSI) (range and sd); range size change between the LGM and the present. | Environmental layers; SDMs | Variables representing long-term climatic stability and historical range dynamics, expected to influence population persistence, bottlenecks and the accumulation of genetic diversity through time. |
| Topography | Elevation (range and sd) | Environmental layers | Variables representing topographic heterogeneity, expected to influence population isolation, environmental heterogeneity and the availability of climatic refugia. |
| Land-cover characteristics | Bare or sparse vegetation, built-up areas, cropland, grassland, mangroves, shrubland, tree cover, permanent water bodies and herbaceous wetlands (range and sd) | Environmental layers | Variables representing habitat availability and environmental heterogeneity, expected to influence population persistence, dispersal opportunities and effective population size. |
| Species geographic characteristics | Geographic range size | IUCN distribution polygons | Variable representing the spatial extent of species distributions, expected to be positively associated with population size, connectivity and demographic stability |
| Life-history traits | Reptiles: microhabitat, habitat type, maximum body mass, maximum body length, maximum female body length, reproductive mode, mean number of offspring and clutch size range.<br>Amphibians: body size and reproductive output. | Trait databases | Species-specific ecological and life-history characteristics expected to influence demographic resilience, effective population size and genetic drift. |
| Evolutionary history | Taxonomic order; taxonomic family | Taxonomic databases | Taxonomic variables representing shared evolutionary history and conserved biological characteristics that may influence genetic diversity. |
| Conservation status | IUCN Red List conservation category | IUCN Red List | Species conservation category representing long-term demographic decline and population fragmentation, which may reduce genetic diversity. |
| Sampling bias<br>(control<br>variables) | Mean number of genetic<br>samples per mitochondrial<br>marker; range<br>representativeness | Genetic dataset | Control variables<br>included to account for<br>biases associated with<br>uneven genetic sampling<br>effort and spatial<br>coverage among species. |

The compilation of predictor variables first required assembling a curated set of species occurrence records, which was subsequently used to extract environmental variables and derive species distribution model (SDM)-based predictors. Because several ecological processes cannot be adequately represented by a single variable, multiple predictors were compiled for each hypothesis to capture complementary aspects of the underlying mechanisms. Some variables were obtained directly from existing spatial layers, species range maps, taxonomic databases or trait databases. Other variables were derived indirectly from SDMs fitted using the curated occurrence records, allowing us to characterize species-level climatic niches, niche breadth, marginality and historical range dynamics. The compiled variables were then used to evaluate the relative importance of alternative ecological hypotheses in explaining variation in species-level genetic diversity across amphibians and reptiles.

### 2.4. Occurrence data compilation and curation

To derive predictor variables based on SDMs and provide a common spatial basis for environmental variable extraction, we assembled a comprehensive set of occurrence records for each species by combining georeferenced genetic sampling localities with additional biodiversity occurrence records.

We searched for additional species occurrence records using the occ() function with the species name from the spocc R package (Owens et al., 2024) to maximise the coverage of our sampling for the variable compilation. We extracted occurrence data from the Global Biodiversity Information Facility (GBIF), VertNet and Integrated Digitized Biocollections (iDigBio). These occurrences were combined with the corresponding georeferenced sequence data and cleaned using the clean_coordinates() function from the CoordinateCleaner R package (Zizka et al., 2019). We then clipped the occurrence records, ensuring that those falling largely outside species’ IUCN ranges were removed. For this purpose, we assigned a buffer to increase proportionally to the square root of the range area, from 10 km for the smallest ranges (≤100 km^2^) to 100 km for the largest ranges (≥8,101 km^2^) in increments of 10 km. These occurrences were then thinned to remove occurrence records that shared the same grid cell at ∼2.5 km resolution. The resulting dataset constituted the curated localities used to extract environmental variables and derive SDM-based predictors for each species (hereafter, curated localities).

### 2.5. Water and energy availability, topography and historical climatic stability

To represent the hypotheses related to water and energy availability, topography and historical climatic stability, we compiled climatic, topographic and historical environmental variables from publicly available spatial datasets. To represent water and energy availability (Table 2), we compiled seven bioclimatic variables from CHELSA v2.1. (Karger et al., 2017) at 30-seconds spatial resolution (∼1 km). These comprised annual extremes for temperature (BIO5: mean daily maximum air temperature of the warmest month; BIO6: mean daily minimum air temperature of the coldest month) and precipitation (BIO16: mean monthly precipitation amount of the wettest quarter; BIO17: mean monthly precipitation amount of the driest quarter), both annual temperature and precipitation averages (BIO1: mean annual air temperature; BIO12: annual precipitation amount), and isothermality (BIO3). These climatic factors are well-known to define species’ range limits and reflect climatic tolerances due to their influence on species’ physiology, reproduction and life-history traits (Araújo et al., 2006; Mi et al., 2023).

To represent topographic heterogeneity, we downloaded altitude from Worldclim v.2.1 (Fick & Hijmans, 2017) at 30-seconds spatial resolution. To represent historical climatic stability, we downloaded the Climate Stability Index (CSI) map (Herrando-Moraira et al., 2022), which spans climatic variability from the Pliocene until the present, at 2.5-minutes spatial resolution (∼5 km). All climatic, topographic and climate stability layers were resampled to ∼2.5 km spatial resolution using the resample() function with bilinear interpolation from the raster R package (Hijmans, 2024a). For each species, we extracted the values of each of these variables at the curated localities using the extract() function from the raster R package, and calculated the range and sd for each (Table 2). We chose to use range and sd rather than averages, as they better capture the full spectrum of environmental and topographic conditions, as well as the degree of variation within each species’ range.

To characterize species’ climatic niches (Table 2), we built an ensemble of SDMs (Appendix S1) and derived four complementary climatic niche metrics: temperature niche breadth, precipitation niche breadth, climatic niche hypervolume and climatic niche marginality. Temperature and precipitation niche breadth were quantified using annual climatic extremes because these variables are known to define species’ climatic tolerances and geographic range limits (*e.g.*, Liu et al., 2020; Moreira et al., 2024).

Temperature niche breadth was calculated as the difference between the maximum value of BIO5 and the minimum value of BIO6 across all niche-envelope localities. Precipitation niche breadth was calculated as the difference between the maximum value of BIO16 and the minimum value of BIO17 across all niche-envelope localities. Climatic niche hypervolume was estimated by i) calculating a PCA with the full dataset of all species’ niche-envelope localities (separately for reptiles and amphibians), ii) filtering the PCA dataset per species, and iii) calculating species-specific hypervolume from the first four PCA axes using the function hypervolume_gaussian() from the hypervolume R package (Blonder et al., 2025). Using a common PCA ensured that hypervolume estimates were directly comparable among species while reducing dimensionality and computational demands. Climatic niche marginality was quantified by calculating the Euclidean distance between each species’ centroid and the grand centroid in the multidimensional climatic space defined by the first four principal component axes, using the PCA scores generated for the climatic niche hypervolume analysis.

To represent land-cover characteristics (Table 2), we compiled nine land-cover classes: bare/sparse vegetation, built-up, cropland, grassland, mangroves, shrubland, tree cover, permanent water bodies and herbaceous wetland from ESA WorldCover (Zanaga et al., 2021) at 30-seconds spatial resolution, using the function landcover() from the geodata R package (Hijmans, Barbosa, et al., 2024). Following the same workflow described for the climatic variables, all layers were resampled to ∼2.5 km spatial resolution, values were extracted at the curated localities, and the range and standard deviation of each land-cover class were calculated for every species. To represent species’ geographic characteristics (Table 2) we calculated the range size (km^2^) of each species’ IUCN distribution polygon using the function expanse() from the terra R package (Hijmans, 2024b). To represent historical range dynamics (Table 2), we quantified changes in species’ distributions between the LGM and the present using the function BIOMOD_RangeSize() from the biomod2 R package (Thuiller et al., 2024), which estimates the percentage of pixels predicted to change between binary projections of the ensembled SDMs of both time steps. Positive values indicate range contraction, negative values indicate range expansion, and values equal to zero indicate no predicted change in range size.

To represent life-history traits (Table 2), we compiled species-specific ecological and reproductive traits. For amphibians, the compiled traits included habitat, diel, body mass, minimum and maximum age at maturity, body size, minimum and maximum size at maturity, maximum longevity, minimum and maximum litter size, reproductive output, minimum and maximum offspring size, and breeding strategy, which were downloaded from AmphiBIO (Oliveira et al., 2017). For reptiles, the compiled traits included main biogeographic region, microhabitat, habitat type, active time, maximum longevity, maximum body mass, maximum length, maximum female length, maximum male length, maximum juvenile length, hatching/neonate mass, reproductive mode, mean number of offspring/eggs per litter/clutch, smallest and largest clutch size, and number of litters/clutches produced per year, which were downloaded from ReptTraits (Oskyrko et al., 2024). We calculated clutch size ranges from the largest and smallest clutch sizes of every species. Species lacking data for all ecological traits were excluded from subsequent analyses. We also removed traits with more than 40% missing data and, for the remaining traits, we imputed missing data using the mice() function from the mice R package (van Buuren & Groothuis-Oudshoorn, 2011). We conducted the imputation using 500 iterations, including species taxonomy (*i.e.*, family) as a predictor in the imputation. To improve the accuracy of trait data imputation, we performed the imputation step using the full ecological traits datasets, AmphiBIO and ReptTraits, which include data for approximately 6,500 and 12,000 species, respectively. This allowed us to leverage broader taxonomic information, which is known to enhance imputation reliability. After completing the imputation, we extracted the relevant imputed trait values for our target species. Taxonomy (order and family) and conservation status were downloaded from IUCN (IUCN, 2022).

To account for potential biases associated with uneven genetic sampling among species (Table 2), we included two control variables representing genetic sampling effort and the spatial representativeness of genetic sampling. Genetic sampling effort was quantified as the mean number of sequences per mitochondrial marker, whereas spatial representativeness was estimated using a range representativeness metric that quantifies how well the sampled localities cover each species’ geographic distribution (Figure S1). We estimated range observed representativeness using the function urap.proportion.held() from the raptr R package (Hanson et al., 2018), which calculates the proportion of total variance explained by the observed sample points in comparison to a uniform distribution of points across all the species’ range. Next, we calculated expected representativeness by replacing the observed points by an evenly distribution of the same number of points across the species’ range. This was achieved by performing k-means clustering to classify the raster centroids into a number of clusters equal to the sample size. The centroids of these clusters, calculated using Voronoi polygons in the dismo R package (Hijmans, Phillips, et al., 2024), were treated as the evenly sampled points. We then repeated the procedure used to calculate the observed representativeness. Finally, we computed the relative representativeness as the ratio of observed to expected representativeness.

### 2.6. Identification of correlates of genetic diversity

To identify the main correlates of species-level intraspecific genetic diversity, we modelled mean nucleotide diversity as the response variable using the environmental, geographic, biological and sampling variables summarized in Table 2 as predictor variables. These variables represent the ecological hypotheses proposed in Table 1, allowing us to evaluate their relative importance in explaining variation in species-level genetic diversity. To that purpose, we used random forests modelling (Breiman, 2001), an ensemble machine-learning algorithm that combines multiple decision trees to model complex, non-linear relationships while accommodating interactions among predictor variables. Variable importance is determined by measuring the increase in model error when the variable is omitted (Liaw & Wiener, 2002).

Random forests models were implemented using the caret R package (Kuhn, 2008). Prior to modelling, we computed pairwise Pearson correlations (r) between all predictor variables using the cor() function from the stats R package. To minimise multicollinearity, upon testing different Pearson’s r thresholds (0.75; 0.80; 0.85; 0.90), we removed predictors with a Pearson’s r greater than 0.90 given that it yielded the best model performance. Additionally, we used recursive feature elimination using the function rfe() from the caret R package (Kuhn, 2008). We split our data into training (90%) and testing (10%) and used 2,000 trees and a 5-fold cross validation to train our analyses. Variable importance was estimated using the importance() function from the randomForest R package (Liaw & Wiener, 2002) and averaged across 50 random seeds to obtain a consensus ranking. We assessed model performance using root mean squared error (RMSE).

To better understand and visualise the relationship between the predictor variables and genetic diversity, we divided mean nucleotide diversity into four quartiles depicting low, medium low, medium high and high genetic diversity, using the quantile() function from the stats R package. We then compared the five most important correlates between the low and high genetic diversity classes using *t*-tests and visualized these relationships with boxplots. We focused on the high and low diversity classes, as they represent the most extreme cases, providing clearer insights into the relationships between predictor variables and species-level intraspecific genetic diversity.

## 3. Results

A total of 365 amphibian and 359 reptile species occurring within our six study regions fulfilled the initial inclusion criteria of presenting at least one mitochondrial gene containing ten or more sequences. A total of 89,589 mitochondrial sequences corresponding to these taxa were assembled into a genetic dataset. Only 10.3% (n=9,222) of these sequences had associated geographic coordinates in NCBI. We were able to increase the georeferenced data to 43.8% (n=39,248) by retrieving geographical coordinates from locality descriptors of associated metadata, and to 87.4% (n=78,269; complete dataset) by manually compiling geographical coordinates and/or locality data from publications (see Figure S2-S3). The final list of taxa was reduced to 247 amphibian and 295 reptile species after further removing species without at least three georeferenced sequences (see Dataset S1).

### 3.1. Genetic diversity across regions

Mean scaled nucleotide diversity was significantly different among study regions in both amphibians (Figure 2a, p = 0.00125) and reptiles (Figure 2b, p = 0.000132). The highest values were recovered in the Brazilian AF (0.639 ± 0.199) and Madagascar (0.612 ± 0.233) for amphibians, and in AMT (0.524 ± 0.236) for reptiles.

**Figure 2.**
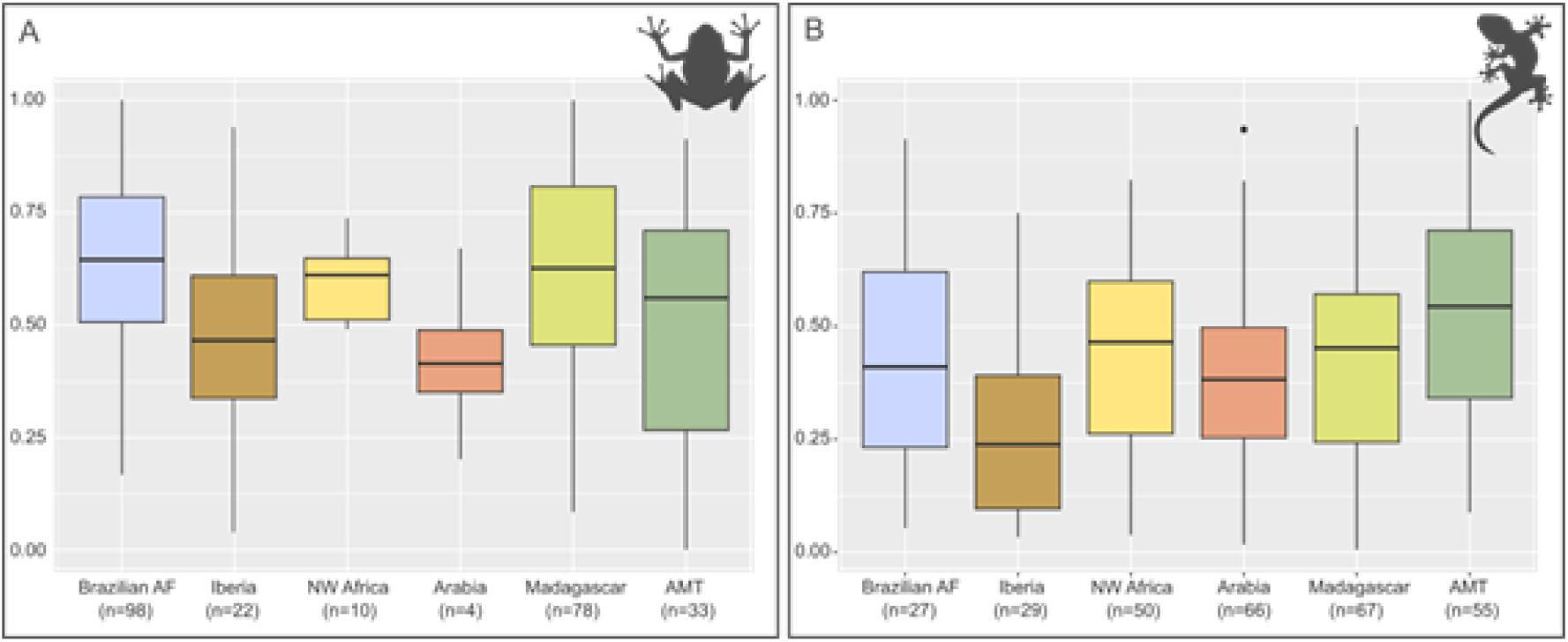
(a) Boxplots of amphibian mean scaled nucleotide diversity per study region. (b) Boxplots of reptile mean scaled nucleotide diversity per study region. Each box–whisker plot indicates the median (bold lines) and the interquartile range (boxes).

### 3.2. Compilation and selection of predictor variables

A total of 49 and 55 environmental, geographic, biological and control variables were compiled for amphibians and reptiles, respectively. Following the removal of highly correlated variables and recursive feature elimination, the final random forests models included 43 variables for amphibians and 52 for reptiles (Table 2; Table S1).

### 3.3. Correlates of genetic diversity

For amphibians, the top-five most important variables in the random forest regression (RMSE = 0.2086, R^2^ = 0.1895) were body size, range and sd of altitude, and range and sd of the mean monthly precipitation amount of the driest quarter—BIO17 (Figure 3a). All top-five predictors were significantly different in species estimated to contain high and low intraspecific diversity (Figure 3b). Furthermore, higher genetic diversity was associated with higher values of all top-five predictors, except in body size, where higher body size was associated with lower genetic diversity.

**Figure 3.**
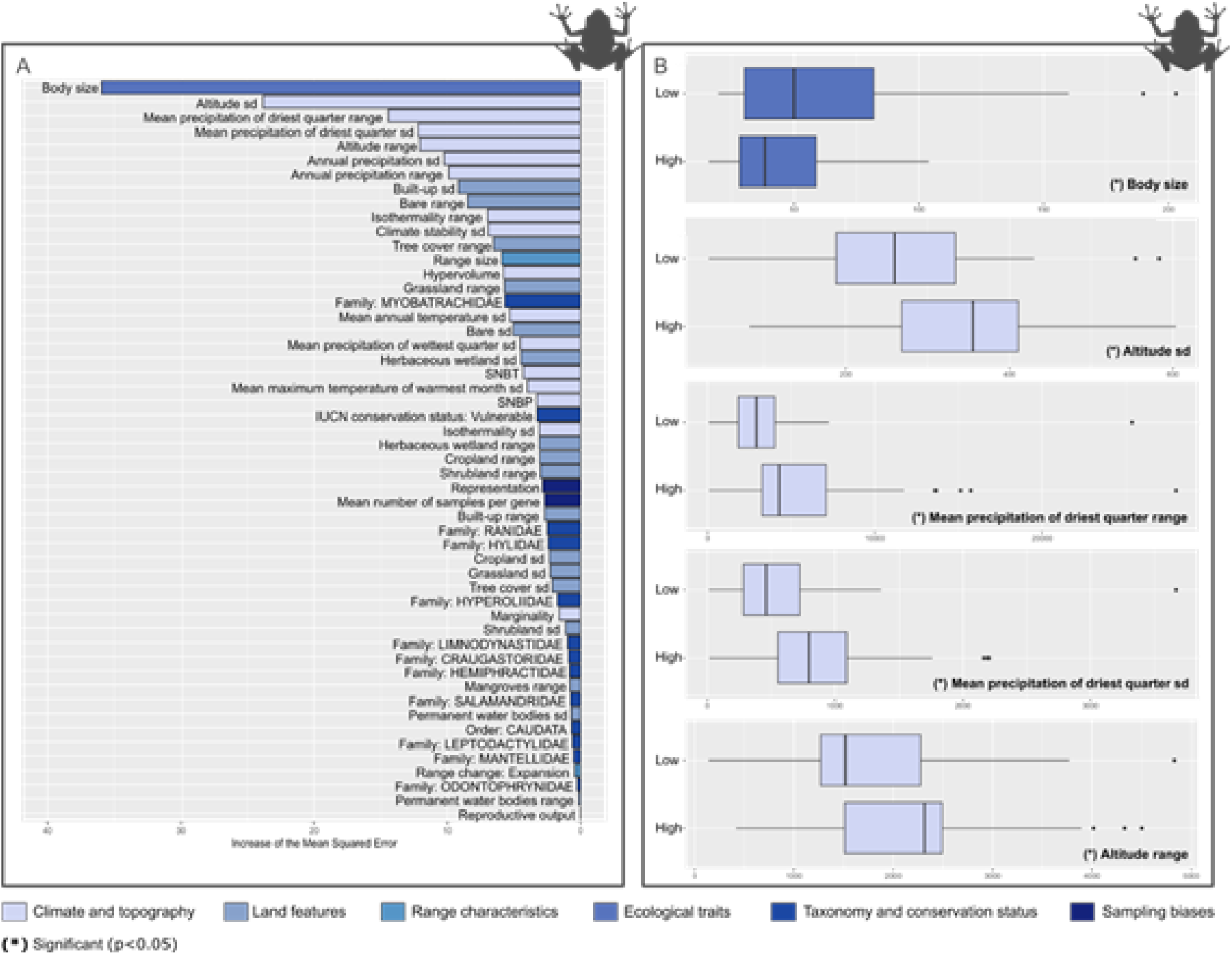
Important correlates of genetic diversity in amphibians. (a) Variable importance ranking according to the consensus random forest regression. (b) Boxplots representing values of the five most important variables according to the consensus random forest regression.

For reptiles, the top-five most important variables in the random forest regression (RMSE = 0.2054, R^2^ = 0.2063) were maximum body mass, range of the mean daily minimum air temperature of the coldest month—BIO6, mean number of offspring, clutch size’s range, and maximum length (Figure 4a). Out of them, only the mean number of offspring and maximum length showed significant differences between species with high and low estimated intraspecific diversity (Figure 4b). Higher genetic diversity was associated with lower values of both variables, showing significant differences.

**Figure 4.**
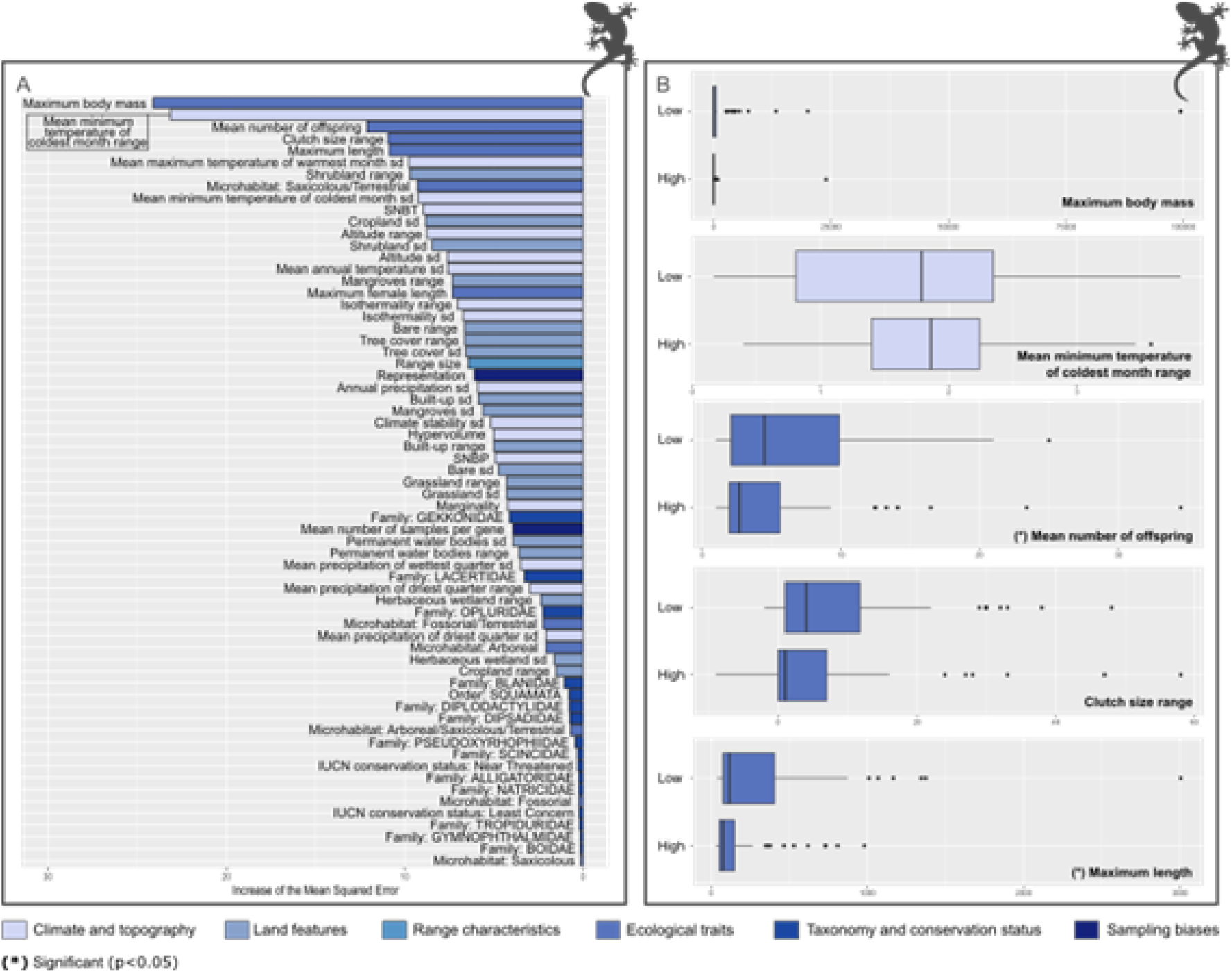
Important correlates of genetic diversity in reptiles. (a) Variable importance ranking according to the consensus random forest regression. (b) Boxplots representing values of the five most important variables according to the consensus random forest regression.

## 4. Discussion

### 4.1. Regional patterns of genetic diversity

We found significant differences in herptile genetic diversity among the six study regions. Tropical regions presented overall higher genetic diversity, as recovered for the Brazilian AF, Madagascar and AMT, a pattern consistent with the extensive discussion of latitudinal gradients in the literature (*e.g.,* Fonseca et al., 2023; Manel et al., 2020; Miraldo et al., 2016; Theodoridis et al., 2020). Tropical regions usually exhibit higher temperature, precipitation and climate stability, all of which may contribute to the maintenance of large and spatially structured populations over evolutionary time. However, the regional differences recovered here may also be influenced by sampling biases, such as the higher number of samples in tropical regions. Taxonomic practice, including recent revisions (*e.g.,* Madagascar) and more generally taxonomic inflation, can further affect estimates of diversity, as the splitting or lumping of geographically structured lineages changes the amount of variation represented within individual species. Additionally, the region of species occurrence was one of the few variables removed during the recursive feature elimination analysis because of its limited explanatory power. This may suggest that the broad geographic regions are not direct determinants of genetic diversity; rather, topographic and environmental conditions, along with the functional traits of species occurring within each region, likely play a more important role in shaping species’ genetic diversity.

### 4.2. Correlates of genetic diversity

The set of environmental, geographic, biological and control variables tested in the random forests analysis explained about 20% of the species-level intraspecific genetic diversity observed. The level of explanatory power is consistent with that reported in other macrogenetic studies targeting different taxonomic groups (*e.g.*, Amador et al., 2024; Barrow et al., 2021; Manel et al., 2020) and reflects the inherently multifactorial nature of species-level genetic diversity. Contemporary genetic diversity emerges from interacting environmental, demographic and evolutionary processes and life-history traits operating across multiple spatial and temporal scales. Importantly, ecological hypotheses are not mutually exclusive, but instead represent complementary mechanisms through which mutation, selection, genetic drift and gene flow shape genetic diversity. Overall, our results provide the strongest support for the hypotheses related to life-history traits, topographic heterogeneity and water-energy availability, whereas comparatively little support was found for the hypothesis that long-term climatic stability is a major determinant of contemporary species-level genetic diversity. The relative importance of these hypotheses differed between taxonomic groups: environmental hypotheses received stronger support in amphibians, whereas life-history traits consistently emerged as the dominant correlates of genetic diversity in reptiles, suggesting that the processes shaping genetic diversity depend on species physiology and life-history.

Our analyses revealed that life-history traits, particularly body size, for amphibians, and maximum length and body mass, for reptiles, are the most important correlates of intraspecific genetic diversity. Amphibians with larger body sizes and reptiles with a larger maximum length present significantly lower intra-specific genetic diversity compared to smaller-bodied species. These findings are consistent with previous studies across several taxonomic groups (but see (Clark et al., 2023; Doyle et al., 2015)) as well as with studies investigating predictors of cryptic diversity (Decker et al., 2025; Parsons et al., 2022). This may corroborate previous findings suggesting that larger-bodied species tend to present smaller population sizes and, consequently, lower genetic diversity (Brüniche-Olsen et al., 2019; Larkin et al., 2023). Additionally, it has also been suggested that species with larger body sizes tend to present lower rates of molecular evolution (Bromham, 2011) and, in turn, lower genetic diversity.

Our results recovered the mean number of offspring as an important correlate of reptile diversity. Higher values were associated with lower genetic diversity, contrary to expectations of a higher number of offspring leading to a higher effective population size, and consequently, to higher genetic diversity (Romiguier et al., 2014). However, reproductive output does not necessarily translate directly into effective population size, as large numbers of offspring may also imply high juvenile mortality and fluctuations in reproductive success, which can lead to genetic drift and population bottlenecks. Additionally, since the focus is on maximizing reproductive output, high fecundity may be associated with minimal parental care, which can cause high mortality and reduce effective population size, limiting genetic diversity. Furthermore, a high population turnover may also result in a reduced accumulation of mutations and stronger genetic drift, which could lead to lower diversity.

Among the environmental hypotheses, topographic heterogeneity and water-energy availability received the strongest support in amphibians. Specifically, the range and standard deviation of elevation and the range and standard deviation of precipitation during the driest quarter were among the strongest correlates of genetic diversity. The role of precipitation has long been discussed, and recent macrogenetic studies recovered a positive relationship with genetic diversity (Amador et al., 2024). Our results also suggest that species occupying a wider variety of altitude and precipitation conditions have significantly higher genetic diversity. Areas with complex environments often support greater genetic diversity due to the variety of microhabitats that can restrict gene flow, favouring population differentiation and local adaptation, which enhance the generation and maintenance of genetic diversity.

Contrary to our expectations, climate stability was not a major correlate of herptile genetic diversity across our study regions. Similar findings were reported by (De Kort et al., 2021), who also found little evidence that past temperature stability favoured population genetic diversity. One possible explanation for this is that ecological traits related to body size and reproductive output can have more immediate effects on genetic diversity, either by shaping population sizes, gene flow, or even local adaptation. Alternatively, the influence of long-term climatic stability may already be partially captured by current environmental conditions and topographic heterogeneity, reducing its apparent importance in the models.

A limitation inherent to macrogenetic analyses is the heterogeneous geographical and taxonomic distribution of sequence data. Although our compilation substantially increased the proportion of georeferenced sequences, species differed considerably in sampling intensity and spatial coverage. Species-wide genetic diversity may therefore reflect not only variation within local populations but also the extent to which geographically differentiated populations are represented in the available sequences. Additionally, while mtDNA remains the most widely available marker for macrogenetic studies, we acknowledge that the extent to which these data are representative of broader genomic patterns remains unclear (Galtier et al., 2009; Larkin et al., 2023). Mitochondrial DNA is inherited maternally, constitutes a small part of the genome and does not undergo recombination, which may limit its ability to capture the full spectrum of genetic variation within populations (Galtier et al., 2009). Thus, while mitochondrial data provides a valuable proxy for large-scale genetic assessments, the correlates of mitochondrial diversity examined here may differ from those shaping genome-wide patterns of variation.

## Acknowledgements

This work was funded by Fundação para a Ciência e Tecnologia, I.P, via national funds, under the scope of project PLACES - "PLAnning for the Conservation of life on Earth in the light of evolution and Sustainability" (DOI 10.54499/2022.04598.CEECIND/CP1730/CT0006).

MJP was supported by Fundação para a Ciência e Tecnologia through a PhD Grant (SFRH/BD/08664/2020). SBC was supported by Fundação para a Ciência e Tecnologia through an individual scientific employment program contract (CEECIND/CP1730/CT0006).

## References

Adams, R. I., & Hadly, E. A. (2013). Genetic diversity within vertebrate species is greater at lower latitudes. Evolutionary Ecology, 27(1), 133–143. 10.1007/s10682-012-9587-x

Allen, A. P., Gillooly, J. F., Savage, V. M., & Brown, J. H. (2006). Kinetic effects of temperature on rates of genetic divergence and speciation. Proceedings of the National Academy of Sciences, 103(24), 9130–9135. 10.1073/pnas.0603587103

Amador, L., Arroyo-Torres, I., & Barrow, L. N. (2024). Machine learning and phylogenetic models identify predictors of genetic variation in Neotropical amphibians. Journal of Biogeography, 51(5), 909–923. 10.1111/jbi.14795

Andrello, M., D’Aloia, C., Dalongeville, A., Escalante, M. A., Guerrero, J., Perrier, C., Torres-Florez, J. P., Xuereb, A., & Manel, S. (2022). Evolving spatial conservation prioritization with intraspecific genetic data. Trends in Ecology & Evolution, 37(6), 553–564. 10.1016/j.tree.2022.03.003

Araújo, M. B., Thuiller, W., & Pearson, R. G. (2006). Climate warming and the decline of amphibians and reptiles in Europe. Journal of Biogeography, 33(10), 1712–1728. 10.1111/j.1365-2699.2006.01482.x

Barrow, L. N., Masiero da Fonseca, E., Thompson, C. E. P., & Carstens, B. C. (2021). Predicting amphibian intraspecific diversity with machine learning: Challenges and prospects for integrating traits, geography, and genetic data. Molecular Ecology Resources, 21(8), 2818–2831. 10.1111/1755-0998.13303

Blanchet, S., Prunier, J. G., & Kort, H. D. (2017). Time to Go Bigger: Emerging Patterns in Macrogenetics. Trends in Genetics, 33(9), 579–580. 10.1016/j.tig.2017.06.007

Blonder, B., Morrow, C. B., Brown, S., Butruille, G., Chen, D., Laini, A., Harris, D. J., & Violet, C. (2025). hypervolume: High Dimensional Geometry, Set Operations, Projection, and Inference Using Kernel Density Estimation, Support Vector Machines, and Convex Hulls (Version R package version 3.1.5) [Computer software]. https://CRAN.R-project.org/package=hypervolume

Breiman, L. (2001). Random Forests. Machine Learning, 45(1), 5–32. 10.1023/A:1010933404324

Bromham, L. (2011). The genome as a life-history character: Why rate of molecular evolution varies between mammal species. Philosophical Transactions of the Royal Society B: Biological Sciences, 366(1577), 2503–2513. 10.1098/rstb.2011.0014

Brüniche-Olsen, A., Kellner, K. F., & DeWoody, J. A. (2019). Island area, body size and demographic history shape genomic diversity in Darwin’s finches and related tanagers. Molecular Ecology, 28(22), 4914–4925. 10.1111/mec.15266

Clark, A., Koc, G., Eyre-Walker, Y., & Eyre-Walker, A. (2023). What Determines Levels of Mitochondrial Genetic Diversity in Birds? Genome Biology and Evolution, 15(5), evad064. 10.1093/gbe/evad064

Convention on Biological Diversity. (2022). Kunming-Montreal Global Biodiversity Framework. CBD. https://www.cbd.int/gbf

De Kort, H., Prunier, J. G., Ducatez, S., Honnay, O., Baguette, M., Stevens, V. M., & Blanchet, S. (2021). Life history, climate and biogeography interactively affect worldwide genetic diversity of plant and animal populations. Nature Communications, 12(1), 516. 10.1038/s41467-021-20958-2

Decker, S. K., Provost, K. L., & Carstens, B. C. (2025). Bats of a feather: Range characteristics and wing morphology predict phylogeographic breaks in volant vertebrates. Frontiers of Biogeography, 18, e139911. 10.21425/fob.18.139911

Dinerstein, E., Olson, D., Joshi, A., Vynne, C., Burgess, N. D., Wikramanayake, E., Hahn, N., Palminteri, S., Hedao, P., Noss, R., Hansen, M., Locke, H., Ellis, E. C., Jones, B., Barber, C. V., Hayes, R., Kormos, C., Martin, V., Crist, E.,… Saleem, M. (2017). An Ecoregion-Based Approach to Protecting Half the Terrestrial Realm. BioScience, 67(6), 534–545. 10.1093/biosci/bix014

Doyle, J. M., Hacking, C. C., Willoughby, J. R., Sundaram, M., & DeWoody, J. A. (2015). Mammalian genetic diversity as a function of habitat, body size, trophic class, and conservation status. Journal of Mammalogy, 96(3), 564–572. 10.1093/jmammal/gyv061

Fick, S. E., & Hijmans, R. J. (2017). WorldClim 2: New 1-km spatial resolution climate surfaces for global land areas. International Journal of Climatology, 37(12), 4302–4315. 10.1002/joc.5086

Foden, W. B., Butchart, S. H. M., Stuart, S. N., Vié, J.-C., Akçakaya, H. R., Angulo, A., DeVantier, L. M., Gutsche, A., Turak, E., Cao, L., Donner, S. D., Katariya, V., Bernard, R., Holland, R. A., Hughes, A. F., O’Hanlon, S. E., Garnett, S. T., Şekercioğlu, Ç. H., & Mace, G. M. (2013). Identifying the World’s Most Climate Change Vulnerable Species: A Systematic Trait-Based Assessment of all Birds, Amphibians and Corals. PLoS ONE, 8(6), e65427. 10.1371/journal.pone.0065427

Fonseca, E. M., Pelletier, T. A., Decker, S. K., Parsons, D. J., & Carstens, B. C. (2023). Pleistocene glaciations caused the latitudinal gradient of within-species genetic diversity. Evolution Letters, 7(5), 331–338. 10.1093/evlett/qrad030

Galtier, N., Nabholz, B., Glémin, S., & Hurst, G. D. D. (2009). Mitochondrial DNA as a marker of molecular diversity: A reappraisal. Molecular Ecology, 18(22), 4541–4550. 10.1111/j.1365-294X.2009.04380.x

González-del-Pliego, P., Freckleton, R. P., Edwards, D. P., Koo, M. S., Scheffers, B. R., Pyron, R. A., & Jetz, W. (2019). Phylogenetic and Trait-Based Prediction of Extinction Risk for Data-Deficient Amphibians. Current Biology, 29(9), 1557–1563.e3. 10.1016/j.cub.2019.04.005

Hanson, J. O., Rhodes, J. R., Possingham, H. P., & Fuller, R. A. (2018). raptr: Representative and adequate prioritization toolkit in R. Methods in Ecology and Evolution, 9(2), 320–330. 10.1111/2041-210X.12862

Herrando-Moraira, S., Nualart, N., Galbany-Casals, M., Garcia-Jacas, N., Ohashi, H., Matsui, T., Susanna, A., Tang, C. Q., & López-Pujol, J. (2022). Climate Stability Index maps, a global high resolution cartography of climate stability from Pliocene to 2100. Scientific Data, 9(1), 48. 10.1038/s41597-022-01144-5

Hewitt, G. (2000). The genetic legacy of the Quaternary ice ages. Nature, 405(6789), 907–913. 10.1038/35016000

Hijmans, R. J. (2024a). raster: Geographic Data Analysis and Modeling (Version R package version 3.6-30) [Computer software]. https://CRAN.R-project.org/package=raster

Hijmans, R. J. (2024b). terra: Spatial Data Analysis (Version R package version 1.7-83) [Computer software]. https://CRAN.R-project.org/package=terra

Hijmans, R. J., Barbosa, M., Ghosh, A., & Mandel, A. (2024). geodata: Download Geographic Data (Version R package version 0.6-2) [Computer software]. https://CRAN.R-project.org/package=geodata

Hijmans, R. J., Phillips, S., Leathwick, J., & Elith, J. (2024). dismo: Species Distribution Modeling (Version R package version 1.3-16) [Computer software]. https://CRAN.R-project.org/package=dismo

IUCN. (2022). The IUCN Red List of Threatened Species. https://www.iucnredlist.org/

Kahle, D., & Wickham, H. (2013). ggmap: Spatial Visualization with ggplot2. The R Journal, 5(1), 144–161.

Karger, D. N., Conrad, O., Böhner, J., Kawohl, T., Krefit, H., Soria-Auza, R. W., Zimmermann, N. E., Linder, H. P., & Kessler, M. (2017). Climatologies at high resolution for the earth’s land surface areas. Scientific Data, 4(1), 170122. 10.1038/sdata.2017.122

Kuhn, M. (2008). Building Predictive Models in R Using the caret Package. Journal of Statistical Software, 28(5), 1–26. 10.18637/jss.v028.i05

Laikre, L., Hoban, S., Bruford, M. W., Segelbacher, G., Allentoft, F. W., Gajardo, G., Rodríguez, A. G., Hedrick, P. W., Heuertz, M., Hohenlohe, P. A., Jaffé, R., Johannesson, K., Liggins, L., MacDonald, A. J., OrozcoterWengel, P., Reusch, T. B. H., Rodríguez-Correa, H., Russo, I.-R. M., Ryman, N., & Vernesi, C. (2020). Post-2020 goals overlook genetic diversity. Science, 367(6482), 1083–1085. 10.1126/science.abb2046

Langin, K. M., Sillett, T. S., Funk, W. C., Morrison, S. A., & Ghalambor, C. K. (2017). Partial support for the central–marginal hypothesis within a population: Reduced genetic diversity but not increased differentiation at the range edge of an island endemic bird. Heredity, 119(1), 8–15. 10.1038/hdy.2017.10

Larkin, I. E., Myers, E. A., Carstens, B. C., & Barrow, L. N. (2023). Predictors of genomic diversity within North American squamates. Journal of Heredity, 114(2), 131–142. 10.1093/jhered/esad001

Lawrence, E. R., Pedersen, E. J., & Fraser, D. J. (2023). Macrogenetics reveals multifaceted influences of environmental variation on vertebrate population genetic diversity across the Americas. Molecular Ecology, 32(16), 4557–4569. 10.1111/mec.17059

Leigh, D. M., Van Rees, C. B., Millette, K. L., Breed, M. F., Schmidt, C., Bertola, L. D., Hand, B. K., Hunter, M. E., Jensen, E. L., Kershaw, F., Liggins, L., Luikart, G., Manel, S., Mergeay, J., Miller, J. M., Segelbacher, G., Hoban, S., & Paz-Vinas, I. (2021). Opportunities and challenges of macrogenetic studies. Nature Reviews Genetics, 22(12), 791–807. 10.1038/s41576-021-00394-0

Liaw, A., & Wiener, M. (2002). Classification and Regression by randomForest. R News, 2(3), 18–22.

Liu, H., Ye, Q., & Wiens, J. J. (2020). Climatic-niche evolution follows similar rules in plants and animals. Nature Ecology & Evolution, 4(5), 753–763. 10.1038/s41559-020-1158-x

Liz, A. V., Rödder, D., Gonçalves, D. V., Velo-Antón, G., Fonseca, M. M., Geniez, P., Crochet, P.-A., & Brito, J. C. (2021). The role of Sahara highlands in the diversification and desert colonization of the Bosc’s fringe-toed lizard. Journal of Biogeography, 48(11), 2891–2906. 10.1111/jbi.14250

Manel, S., Guerin, P.-E., Mouillot, D., Blanchet, S., Velez, L., Albouy, C., & Pellissier, L. (2020). Global determinants of freshwater and marine fish genetic diversity. Nature Communications, 11(1), 692. 10.1038/s41467-020-14409-7

Mi, C., Ma, L., Yang, M., Li, X., Meiri, S., Roll, U., Oskyrko, O., Pincheira-Donoso, D., Harvey, L. P., Jablonski, D., Safaei-Mahroo, B., Ghaffari, H., Smid, J., Jarvie, S., Kimani, R. M., Masroor, R., Kazemi, S. M., Nneji, L. M., Fokoua, A. M. T.,… Du, W. (2023). Global Protected Areas as refuges for amphibians and reptiles under climate change. Nature Communications, 14(1), 1389. 10.1038/s41467-023-36987-y

Miraldo, A., Li, S., Borregaard, M. K., Flórez-Rodríguez, A., Gopalakrishnan, S., Rizvanovic, M., Wang, Z., Rahbek, C., Marske, K. A., & Nogués-Bravo, D. (2016). An Anthropocene map of genetic diversity. Science, 353(6307), 1532–1535. 10.1126/science.aaf4381

Moreira, M. O., Wiens, J. J., Fonseca, C., & Rojas, D. (2024). Climatic-niche breadth, niche position, and speciation in lizards and snakes. Journal of Biogeography, 51(6), 969–981. 10.1111/jbi.14802

Nei, M., & Li, W. H. (1979). Mathematical model for studying genetic variation in terms of restriction endonucleases. Proceedings of the National Academy of Sciences, 76(10), 5269–5273. 10.1073/pnas.76.10.5269

Ochoa-Ochoa, L. M., Mejía-Domínguez, N. R., Velasco, J. A., Dimitrov, D., & Marske, K. A. (2020). Dimensions of amphibian alpha diversity in the New World. Journal of Biogeography, 47(11), 2293–2302. 10.1111/jbi.13948

Ohsawa, T., & Ide, Y. (2008). Global patterns of genetic variation in plant species along vertical and horizontal gradients on mountains. Global Ecology and Biogeography, 17(2), 152–163. 10.1111/j.1466-8238.2007.00357.x

Oliveira, B. F., São-Pedro, V. A., Santos-Barrera, G., Penone, C., & Costa, G. C. (2017). AmphiBIO, a global database for amphibian ecological traits. Scientific Data, 4(1), 170123. 10.1038/sdata.2017.123

Oskyrko, O., Mi, C., Meiri, S., & Du, W. (2024). ReptTraits: A comprehensive dataset of ecological traits in reptiles. Scientific Data, 11(1), 243. 10.1038/s41597-024-03079-5

Owens, H., Barve, V., & Chamberlain, S. (2024). spocc: Interface to Species Occurrence Data Sources (Version R package version 1.2.3) [Computer software]. https://CRAN.R-project.org/package=spocc

Paradis, E. (2010). pegas: An R package for population genetics with an integrated–modular approach. Bioinformatics, 26(3), 419–420. 10.1093/bioinformatics/btp696

Parsons, D. J., Pelletier, T. A., Wieringa, J. G., Duckett, D. J., & Carstens, B. C. (2022). Analysis of biodiversity data suggests that mammal species are hidden in predictable places. Proceedings of the National Academy of Sciences, 119(14), e2103400119. 10.1073/pnas.2103400119

Paz-Vinas, I., Jensen, E. L., Bertola, L. D., Breed, M. F., Hand, B. K., Hunter, M. E., Kershaw, F., Leigh, D. M., Luikart, G., Mergeay, J., Miller, J. M., Van Rees, C. B., Segelbacher, G., & Hoban, S. (2021). Macrogenetic studies must not ignore limitations of genetic markers and scale. Ecology Letters, 24(6), 1282–1284. 10.1111/ele.13732

Pelletier, T. A., Parsons, D. J., Decker, S. K., Crouch, S., Franz, E., Ohrstrom, J., & Carstens, B. C. (2022). phylogatR: Phylogeographic data aggregation and repurposing. Molecular Ecology Resources, 22(8), 2830–2842. 10.1111/1755-0998.13673

Peng, S., Shrestha, N., Luo, Y., Li, Y., Cai, H., Qin, H., Ma, K., & Wang, Z. (2023). Incorporating global change reveals extinction risk beyond the current Red List. Current Biology, 33(17), 3669–3678.e4. 10.1016/j.cub.2023.07.047

Portik, D. M., & Wiens, J. J. (2020). SuperCRUNCH: A bioinformatics toolkit for creating and manipulating supermatrices and other large phylogenetic datasets. Methods in Ecology and Evolution, 11(6), 763–772. 10.1111/2041-210X.13392

R Core Team. (2024). R: A Language and Environment for Statistical Computing. [Computer software]. https://www.R-project.org/

Rahbek, C., Borregaard, M. K., Colwell, R. K., Dalsgaard, B., Holt, B. G., Morueta-Holme, N., Nogues-Bravo, D., Whittaker, R. J., & Fjeldså, J. (2019). Humboldt’s enigma: What causes global patterns of mountain biodiversity? Science, 365(6458), 1108–1113. 10.1126/science.aax0149

Romiguier, J., Gayral, P., Ballenghien, M., Bernard, A., Cahais, V., Chenuil, A., Chiari, Y., Dernat, R., Duret, L., Faivre, N., Loire, E., Lourenco, J. M., Nabholz, B., Roux, C., Tsagkogeorga, G., Weber, A., Weinert, L. A., Belkhir, K., Bierne, N.,… Galtier, N. (2014). Comparative population genomics in animals uncovers the determinants of genetic diversity. Nature, 515(7526), 261–263. 10.1038/nature13685

Schluter, D., & Pennell, M. W. (2017). Speciation gradients and the distribution of biodiversity. Nature, 546(7656), 48–55. 10.1038/nature22897

Schmidt, C., Hoban, S., & Jetz, W. (2023). Conservation macrogenetics: Harnessing genetic data to meet conservation commitments. Trends in Genetics, 39(11), 816–829. 10.1016/j.tig.2023.08.002

Sgrò, C. M., Lowe, A. J., & Hoffmann, A. A. (2011). Building evolutionary resilience for conserving biodiversity under climate change. Evolutionary Applications, 4(2), 326–337. 10.1111/j.1752-4571.2010.00157.x

Singhal, S., Huang, H., Title, P. O., Donnellan, S. C., Holmes, I., & Rabosky, D. L. (2017). Genetic diversity is largely unpredictable but scales with museum occurrences in a species-rich clade of Australian lizards. Proceedings of the Royal Society B: Biological Sciences, 284(1854), 20162588. 10.1098/rspb.2016.2588

Theodoridis, S., Fordham, D. A., Brown, S. C., Li, S., Rahbek, C., & Nogues-Bravo, D. (2020). Evolutionary history and past climate change shape the distribution of genetic diversity in terrestrial mammals. Nature Communications, 11(1), 2557. 10.1038/s41467-020-16449-5

Thuiller, W., Georges, D., Gueguen, M., Engler, R., Breiner, F., Lafourcade, B., Patin, R., & Blancheteau, H. (2024). biomod2: Ensemble Platform for Species Distribution Modeling (Version R package version 4.2-5-2) [Computer software]. https://CRAN.R-project.org/package=biomod2

van Buuren, S., & Groothuis-Oudshoorn, K. (2011). mice: Multivariate Imputation by Chained Equations in R. Journal of Statistical Software, 45(3), 1–67. 10.18637/jss.v045.i03

Winter, D. J. (2017). rentrez: An R package for the NCBI eUtils API. The R Journal, 9(2), 520–526. 10.7287/peerj.preprints.3179v2

Zanaga, D., Van De Kerchove, R., De Keersmaecker, W., Souverijns, N., Brockmann, C., Quast, R., Wevers, J., Grosu, A., Paccini, A., Vergnaud, S., Cartus, O., Santoro, M., Fritz, S., Georgieva, I., Lesiv, M., Carter, S., Herold, M., Li, L., Tsendbazar, N.-E.,… Arino, O. (2021). ESA WorldCover 10 m 2020 v100 (Version v100) [Dataset]. Zenodo. 10.5281/zenodo.5571936

Zizka, A., Silvestro, D., Andermann, T., Azevedo, J., Duarte Ritter, C., Edler, D., Farooq, H., Herdean, A., Ariza, M., Scharn, R., Svantesson, S., Wengström, N., Zizka, V., & Antonelli, A. (2019). CoordinateCleaner: Standardized cleaning of occurrence records from biological collection databases. Methods in Ecology and Evolution, 10(5), 744–751. 10.1111/2041-210X.13152

