## Supporting Information for "What shapes intraspecific genetic diversity in amphibians and reptiles? A comparative macrogenetic study across six biogeographic regions"

#### Contents

Appendix S1

Figures S1–S3

#### Other materials

Table S1: Variable predictors compiled in this study driving genetic diversity in herptiles.

Dataset S1.xlsx: Amphibian and reptile presence across study areas and available genetic data.

Dataset S2.xlsx: Amphibian and reptile datasets used in the random forest regression analyses.

R scripts

### APPENDIX S1

#### Ecological modelling

In order to estimate climate niche parameters for subsequent analysis (see Methods section 2.5), for each species, we firstly built an ensemble of species distribution models (SDMs) for the climatic niche using the ensemble() function from the *sdm* R package (Naimi and Araújo, 2016). As input, we considered the curated localities and seven bioclimatic variables (BIO5, BIO6, BIO16, BIO17, BIO1, BIO12 and BIO3) downloaded from CHELSA v2.1. (Karger *et al.*, 2017) at 30-seconds spatial resolution (~1 km).

We constrained the SDM calibration area to a 200 km buffer around the curated localities (VanDerWal *et al.*, 2009; Merow *et al.*, 2013; Mi *et al.*, 2023), to ensure we included only the most likely area (Holt *et al.*, 2013) where each species occurs. We randomly generated pseudo-absences within the calibration area, matching the number of occurrence records per species (with a minimum of 1,000 points) (Barbet-Massin *et al.*, 2012; Mi *et al.*, 2023), using the sdmData() function method from the *sdm* R package. Additionally, considering this calibration area, we removed highly correlated variables with a variance inflation factor >10 from the seven initial bioclimatic variables, using the vifstep() function from the *usdm* R package (Naimi *et al.*, 2014). We then fitted and evaluated SDMs using the sdm() function from the *sdm* R package. For this, we considered five commonly used and high performance algorithms (GLM: generalized linear models; GBM: generalized boosted regression models; Maxent: maximum entropy; RF: random forests; SVM: support vector machine), each repeated five times using bootstrapping replication. We evaluated models using true skill statistics (TSS) and the area under the curve (AUC), and discarded those with a TSS < 0.7. We then built the ensemble based on the retained models, weighted by their performance (TSS). Suitable climatic-niche models were converted to presence/absence based on the averaged threshold that maximises the sum of sensitivity and specificity (maxSSS) across the retained models included in the ensemble (Liu *et al.*, 2013). We clipped each binary ensemble model following the approach of Kremen *et al.*, (2008) to limit the potential over-prediction of SDMs. This method is based on a buffered minimum convex polygon (MCP) around the curated localities, producing models that exclude suitable climatic niche largely outside species′ known occurrence (Brown *et al.*, 2014). As above, the buffer was proportional to the area of the MCP, ranging from 10 km to 100 km depending on the size of the species range. Finally, we generated 5,000 random points (or less, when there were not enough suitable grids in the raster) of suitable climatic niche. Together with the curated localities, these coordinates represent the climatic-niche envelope for each species; hereafter, niche-envelope localities. We also hindcasted the SDMs using the ensemble() function, projecting the retained models (TSS > 0.7) to the Last Glacial Maximum (LGM; 22k before present). To represent the LGM, we downloaded the same seven bioclimatic variables used in the SDMs from CHELSA-TraCE21k v2.1. (Karger *et al.*, 2023) at 30-seconds spatial resolution (~1 km).

#### REFERENCES


Barbet-Massin M, Jiguet F, Albert CH, Thuiller W (2012). Selecting pseudo-absences for species distribution models: how, where and how many? *Methods in Ecology and Evolution* **3**: 327–338.

Brown JL, Cameron A, Yoder AD, Vences M (2014). A necessarily complex model to explain the biogeography of the amphibians and reptiles of Madagascar. *Nat Commun* **5**: 5046.

Holt BG, Lessard J-P, Borregaard MK, Fritz SA, Araújo MB, Dimitrov D, *et al.* (2013). An Update of Wallace's Zoogeographic Regions of the World. *Science* **339**: 74–78.

Karger DN, Conrad O, Böhner J, Kawohl T, Kreft H, Soria-Auza RW, *et al.* (2017). Climatologies at high resolution for the earth's land surface areas. *Sci Data* **4**: 170122.

Karger DN, Nobis MP, Normand S, Graham CH, Zimmermann NE (2023). CHELSA-TraCE21k – high-resolution (1 km) downscaled transient temperature and precipitation data since the Last Glacial Maximum. *Climate of the Past* **19**: 439–456.

Kremen C, Cameron A, Moilanen A, Phillips SJ, Thomas CD, Beentje H, *et al.* (2008). Aligning Conservation Priorities Across Taxa in Madagascar with High-Resolution Planning Tools. *Science* **320**: 222–226.

Liu C, White M, Newell G (2013). Selecting thresholds for the prediction of species occurrence with presence-only data. *Journal of Biogeography* **40**: 778–789.

Merow C, Smith MJ, Silander Jr JA (2013). A practical guide to MaxEnt for modeling species′ distributions: what it does, and why inputs and settings matter. *Ecography* **36**: 1058–1069.

Mi C, Ma L, Yang M, Li X, Meiri S, Roll U, *et al.* (2023). Global Protected Areas as refuges for amphibians and reptiles under climate change. *Nat Commun* **14**: 1389.

Naimi B, Araújo MB (2016). sdm: a reproducible and extensible R platform for species distribution modelling. *Ecography* **39**: 368–375.

Naimi B, Hamm NAS, Groen TA, Skidmore AK, Toxopeus AG (2014). Where is positional uncertainty a problem for species distribution modelling? *Ecography* **37**: 191–203.

VanDerWal J, Shoo LP, Graham C, Williams SE (2009). Selecting pseudo-absence data for presence-only distribution modeling: How far should you stray from what you know? *Ecological Modelling* **220**: 589–594.

**Figure S1.** (a) Boxplots of amphibian range representation per study region. (b) Boxplots of reptile range representation per study region. Each box–whisker plot indicates the median (bold lines) and the interquartile range (boxes).

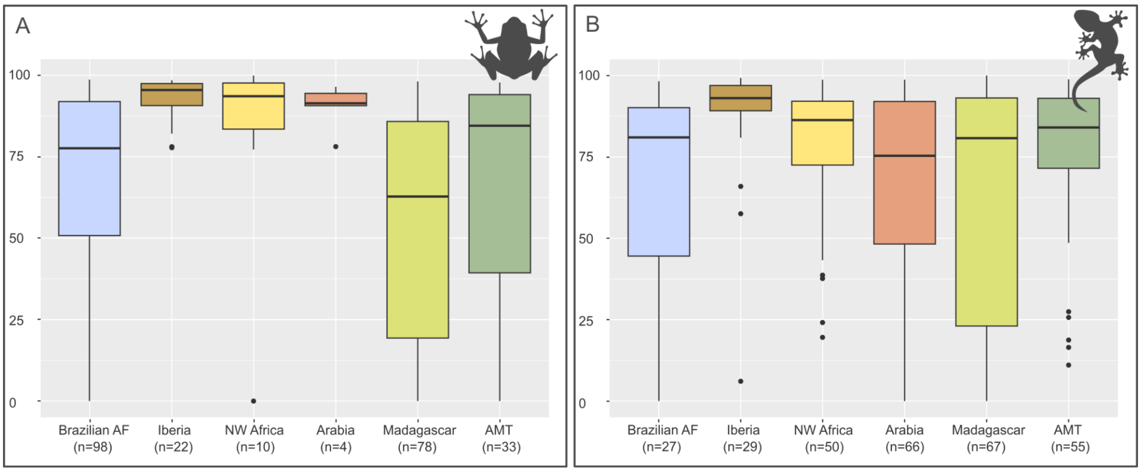

**Figure S2.** Spatial distribution of amphibian mtDNA samples across study region: (a) Brazilian AF, (b) Iberian Peninsula, (c) NW Africa, (d) Arabian Peninsula, (e) Madagascar and (f) AMT.

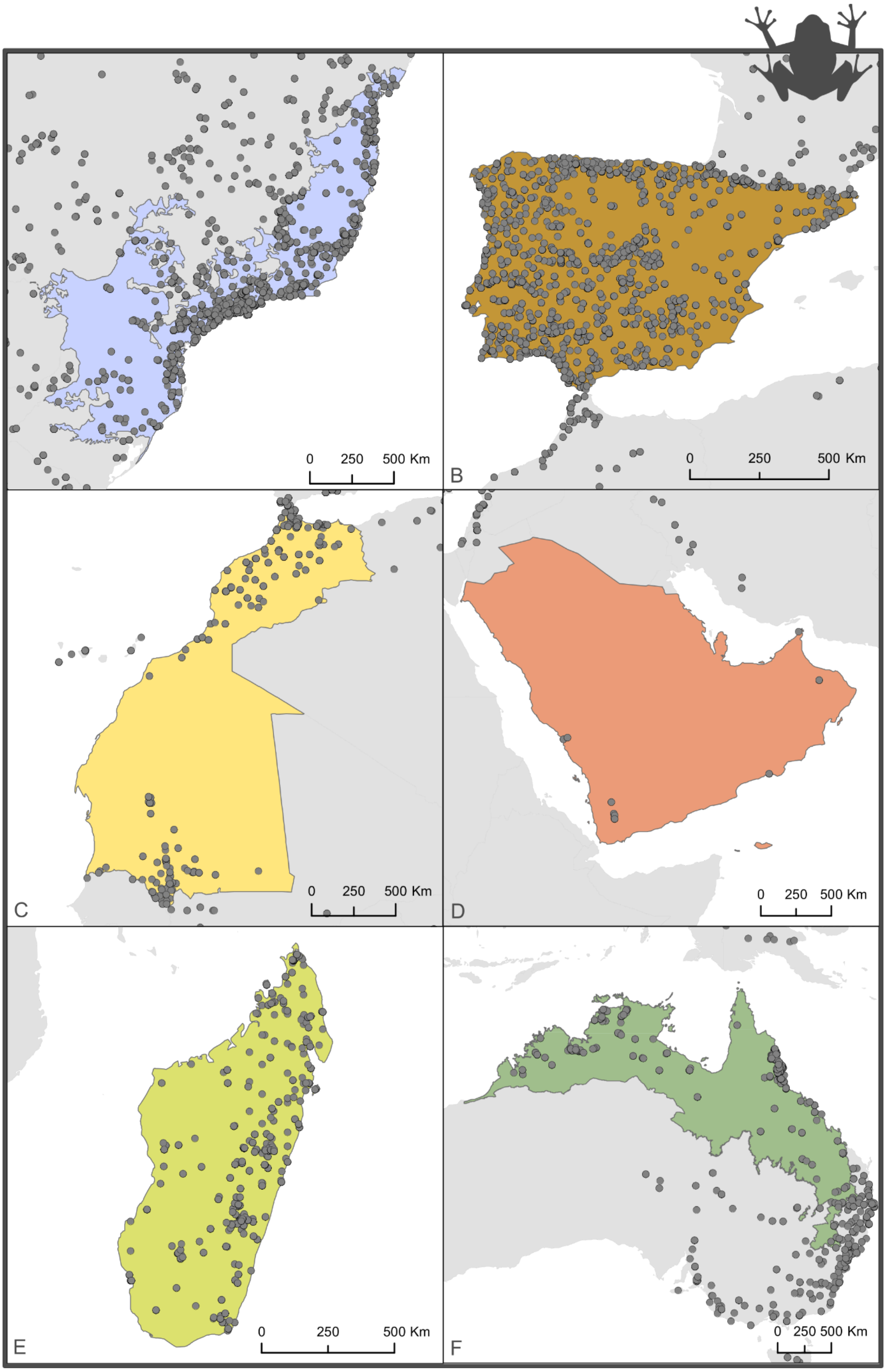

**Figure S3.** Spatial distribution of reptile mtDNA samples across study region: (a) Brazilian AF, (b) Iberian Peninsula, (c) NW Africa, (d) Arabian Peninsula, (e) Madagascar and (f) AMT.

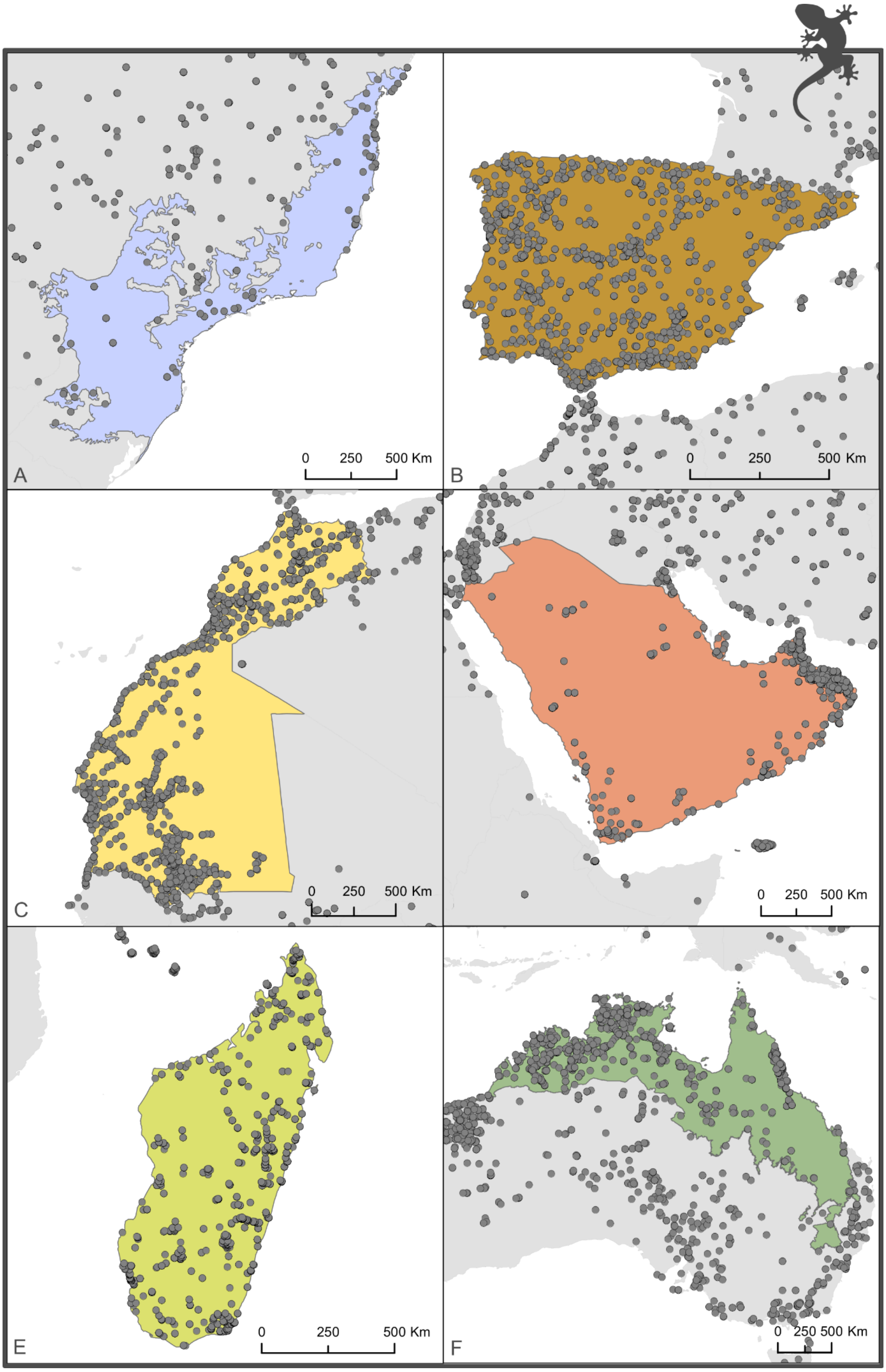
